# Antigen-specific T-cell receptor signatures of cytomegalovirus infection

**DOI:** 10.1101/450882

**Authors:** Alina Huth, Xiaoling Liang, Stefan Krebs, Helmut Blum, Andreas Moosmann

**Affiliations:** DZIF Research Group “Host Control of Viral Latency and Reactivation” (HOCOVLAR), Research Unit Gene Vectors, Helmholtz Zentrum München, Munich, Germany; German Center for Infection Research (DZIF – Deutsches Zentrum für Infektionsforschung), Munich, Germany; HRYZ Biotech Co., Shenzhen, China; Laboratory for Functional Genome Analysis (LAFUGA), Gene Center, Ludwig-Maximilians-Universität München, Munich, Germany

## Abstract

Cytomegalovirus (CMV) is a prevalent human pathogen. The virus cannot be eliminated from the body, but is kept in check by CMV-specific T cells. Patients with an insufficient T-cell response, such as transplant recipients, are at high risk of developing CMV disease. However, the CMV-specific T-cell repertoire is complex, and is not yet clear which T cells protect best against virus reactivation and disease. Here we present a highly resolved characterization of CMV-specific CD8+ T cells based on enrichment by specific peptide stimulation and mRNA sequencing of their T-cell receptor β chains (TCRβ). Our analysis included recently identified T-cell epitopes restricted through HLA-C, whose presentation is resistant to viral immunomodulation, and well-studied HLA-B-restricted epitopes. In 8 healthy virus carriers, we identified a total of 1052 CMV-specific TCRβ chains. HLA-C-restricted, CMV-specific TCRβ clonotypes the *ex vivo* T-cell response, and contributed the highest-frequency clonotype of the entire repertoire in 2 of 8 donors. We analyzed sharing and similarity of CMV-specific TCRβ sequences and identified 63 public or related sequences belonging to 17 public TCRβ families. In our cohort and in an independent cohort of 352 donors, the cumulative frequency of these public TCRβ family members was a highly discriminatory indicator of carrying both CMV infection and the relevant HLA type. Based on these findings, we propose CMV-specific TCRβ signatures as a biomarker for an antiviral T-cell response to identify patients in need of treatment and to guide future development of immunotherapy.

## Introduction

Like other members of the herpesvirus family, human cytomegalovirus (CMV) infects its carriers for life, and prevention of overt disease requires a protective repertoire of virus-specific T cells (1, 2). Persons who lack such a T-cell repertoire, such as patients after allogeneic hematopoietic stem cell transplantation (allo-HSCT), are at risk of reactivating latent CMV infection, and CMV disease remains a threat to their survival and well-being (3). Conventional antiviral chemotherapy has significant adverse effects, is not universally effective, may delay viral reactivation rather than prevent it, and is subverted by viral resistance (3, 4).

Re-establishment of a functional and durable antiviral T-cell repertoire is expected to enable patients to control CMV infection for their lifetime. Transfer of CMV-specific T cells from immunocompetent donors to patients after allo-HSCT has yielded encouraging results over the last three decades (5, 6). However, most of these trials were of small to medium scale and without randomized controls. Preliminary reports on two recent large randomized controlled trials suggest efficacy (5), but complete analysis is not yet available.

Therefore, present capability to clinically exploit the CMV-specific T-cell repertoire is limited. Although the CMV-specific T-cell response has been studied in great detail (7), fundamental questions remain unanswered. No consensus has been reached on which viral antigens and epitopes induce T-cell responses that directly protect against infection, and which T-cell specificities are merely correlated with the presence of other, more effective specificities (7). The challenge of understanding the CMV-specific T-cell repertoire is formidable, since CMV expresses more than 200 viral proteins and has evolved multiple mechanisms to interfere with T-cell recognition by modulating cytokine and chemokine responses, antigen processing, intracellular peptide translocation, stability of HLA molecules, and more (8, 9). Moreover, human CMV shows sequence variation, and some specific T cells may recognize only certain strain-specific epitope variants (10).

A particularly strong CD8+ T-cell response is directed to two viral antigens with contrasting functional roles and kinetics of expression: the transcription factor IE-1 and the structural protein pp65 (11, 12). Earlier data indicated that pp65-specific CD8+ T cells were the most effective in attacking infected cells *in vitro* (13), whereas IE-1-specific T cells were more strongly associated with reduced viral reactivation in patients after transplantation (14, 15) and protective in the murine CMV model (16). Our recent findings suggested that viral immunoevasion is not predominantly guided by the identity of the antigen, but by the identity of the epitope and the HLA class I molecule that presents it (17, 18). For example, we identified an HLA-C-restricted CD8+ T-cell epitope from IE-1 whose presentation is highly resistant to viral immunomodulation. T cells specific for this epitope are of high incidence and frequency in healthy donors (17, 18). These properties are shared (19) by a second CD8+ T-cell epitope restricted through the same HLA (20); this epitope is derived from the rarely studied UL29/28-encoded CMV protein. It is unknown whether there is a causal relationship between escape of certain epitopes from viral immunomodulation and high incidence of epitope-specific T cells, and whether T cells against such epitopes are associated with protection from reactivation. To address such questions, the CMV-specific T-cell response needs to be analyzed and understood in much more detail, using methods that are of sufficient resolution to adequately cover the complexity of the repertoire. High-resolution sequencing of the T-cell receptor (TCR) is such a method.

Most human T cells express a heterodimeric αβ TCR that specifically recognizes the antigenic target, a complex of an HLA molecule and a peptide. Both chains, α and β, have highly variable sequences. The specificity of each αβ T cell is ensured by its expression of only one TCRβ chain and one or, occasionally, two TCRα chains (21). Variability of TCR sequence is produced by recombination in the thymus. In the case of the human TCRβ chain, a V(D)J reading frame is produced by imprecisely joining one of 46 functional V genes, one of two short D genes, and one of 13 J genes, mostly with insertion of template-independent nucleotides between the genes (22, 23). The sequence around these junctions encodes the CDR3, a loop that reaches out to the peptide embedded in the HLA molecule (24). The number of different TCRβ chains in the T-cell repertoire of a human being was estimated to be in the range of millions (25–27) or even hundreds of millions (28); this is only a small fraction of the diversity that is theoretically possible (22, 27). CMV-specific TCR repertoires have been studied in detail before (29–31), but most studies were limited to pp65 and HLA-C-restricted T cells were not included. The advent of massively parallel sequencing of TCR-encoding DNA or mRNA (26, 32) has now made it possible to identify the specificity-defining element of millions of T cells in a sample, and pioneering studies have applied this technique to the analysis of CMV-specific T cells (33–36).

Here we use high-resolution TCRβ sequencing to investigate the repertoire of CMV-specific CD8+ T cells, focusing on previously unstudied HLA-C-restricted T cells that promise to be of high clinical interest. Specific T cells were selectively enriched by peptide-driven *in vitro* expansion; this method is transferable to settings when samples are small or HLA/peptide multimers are not available. We found that T cells specific for HLA-C-restricted CMV epitopes showed exceptional clonal dominance within the overall TCRβ repertoire. Moreover, we identified a set of public and related CMV-specific TCRβ sequences that reliably distinguished persons with or without CMV-specific T-cell immunity, immediately suggesting future application of this method in clinical immunomonitoring.

## Materials and Methods

### Blood donors

Human T cells were derived from anonymous peripheral blood buffy coats purchased from Institut für Transfusionsmedizin, Ulm, Germany. The institutional review board (Ethikkommission bei der LMU München, Project No. 17-455, 16.10.2017) has approved our use of anonymous human material. We did not seek or obtain consent since all material and data were obtained anonymously. Peripheral blood mononuclear cells (PBMCs) were isolated by density centrifugation and cryopreserved until use. Donors were HLA-typed at 4-digit resolution (MVZ, Martinsried, Germany). All donors were positive for HLA-B*07:02 and HLA-C*07:02. CMV status (Supplementary Table 1) was determined by anti-CMV IgG ELISA (Siemens).

### Peptide stimulation assay

The four CMV-derived peptides CRVLCCYVL (CRV, HLA-C*07:02, IE-1), FRCPRRFCF (FRC, HLA-C*07:02, UL28), RPHERNGFTVL (RPH, HLA-B*07:02, pp65), and TPRVTGGGAM (TPR, HLA-B*07:02, pp65) (Supplementary Table 1) were separately used to stimulate and selectively expand virus-specific T cells from CMV-positive donors P01-P08. Cell culture medium was RPMI 1640 (Invitrogen) supplemented with 8% or 10% FCS (Bio-Sell or Invitrogen). Per culture, 25 million PBMCs were suspended in 2 mL cell culture medium containing 5 µg/ml of peptide (JPT, Berlin; ≥70% purity), incubated at 37°C for 1 hour, and washed three times with PBS (PAN Biotech) to remove excess peptide. PBMCs were resuspended in 12.5 ml cell culture medium supplemented with 50 U/ml IL-2 (Proleukin; Novartis) and distributed at 2.5 ml per well to a 12-well plate. The plate was incubated at 37°C and 5% CO_2_. After 6±1 days, the cells of each well were resuspended, distributed to two wells, and 1 ml of fresh culture medium supplemented with IL-2 was added to each well. Cells were harvested at day 10 of culture.

### T-cell stimulation with autologous mini-LCLs

PBMCs from three healthy donors (P01-P03) were infected with empty mini-Epstein-Barr viruses encoding pp65, IE-1, or no CMV protein (37). The resulting transformed B cell lines (mini-lymphoblastoid cell lines, mini-LCLs) were maintained in RPMI 1640 medium supplemented with 8% or 10% FCS. For the stimulation assay, mini-LCLs were γ-irradiated with 50 Gy in a Cs-137 device. Subsequently, 150,000 mini-LCL cells were combined with 6 million PBMCs in 3 mL RPMI/FCS per replicate in a 12-well plate, in four replicates per culture. After 9 days and then every 7 days, the T cells were restimulated: cultures were harvested, washed, counted, and 3 million T cells per well (12-well plate) were coincubated with 1 million irradiated mini-LCLs in medium with 50 U/ml IL-2. At day 30, cultures were harvested to analyze T cells.

### TCRβ library preparation

Where indicated (Supplementary Table 2), CD8+ T cells were enriched from PBMCs or T-cell cultures by magnetic separation with CD8 MicroBeads and MS or LS Columns (Miltenyi Biotech).

Total RNA was extracted from PBMCs, T-cell cultures, or CD8-enriched T cells using the Qiagen RNeasy Kit (Qiagen). 1 µg RNA per sample was reversely transcribed to cDNA with the QuantiTect Reverse Transcription Kit (Qiagen) using a primer designed to target both the Cβ1 and Cβ2 regions of the TCR RNA (5’-GCACC TCCTT CCCAT TCAC-3’). cDNA was amplified in two subsequent PCRs using *Pfu* DNA polymerase on a thermocycler (Biometra T Gradient). Both PCRs were initiated at 95°C for 2 min; cycles consisted of incubation at 95°C (30 sec), 65°C (30 sec), and 72°C (60 sec); final elongation was at 72°C (10 min). The first reaction was a multiplex PCR with 42 distinct forward primers that bind to the Vβ region and cover all possible human TCR Vβ segments, and a reverse primer that anneals to the Cβ1 and Cβ2 regions; the Cβ primer was optimized for this protocol, Vβ binding sites were mostly taken from established protocols (26). Forward (Vβ) and reverse (Cβ) primers carried sequences complementary to the Illumina Read 2 and Read 1 priming sequence, respectively. To enhance the nucleotide diversity of TCRβ reads, facilitate cluster recognition and avoid artefacts, three different forms of the reverse primer were used, with 0, 1 or 2 degenerated nucleotides (N) inserted between the Cβ-binding sequence and the Illumina Read 1 sequence. All primers were used in equimolar amounts (a total of 10 µM; for primer sequences see Supplementary Table 1). The first PCR consisted of only 10 amplification cycles to minimize PCR amplification bias. The second PCR was performed with index primers (NEBNext Multiplex Oligos for Illumina; New England BioLabs), to attach barcodes and the i5 and i7 adapters for cluster generation on the Illumina flow cell (Figure 1B). After each PCR step, the PCR product was purified using Agencourt AMPure XP magnetic beads (Beckman Coulter). Length and quantity of PCR products for sequencing was determined using the Agilent DNA 1000 Kit and the Bioanalyzer 2100 (Agilent).

**Figure 1.**
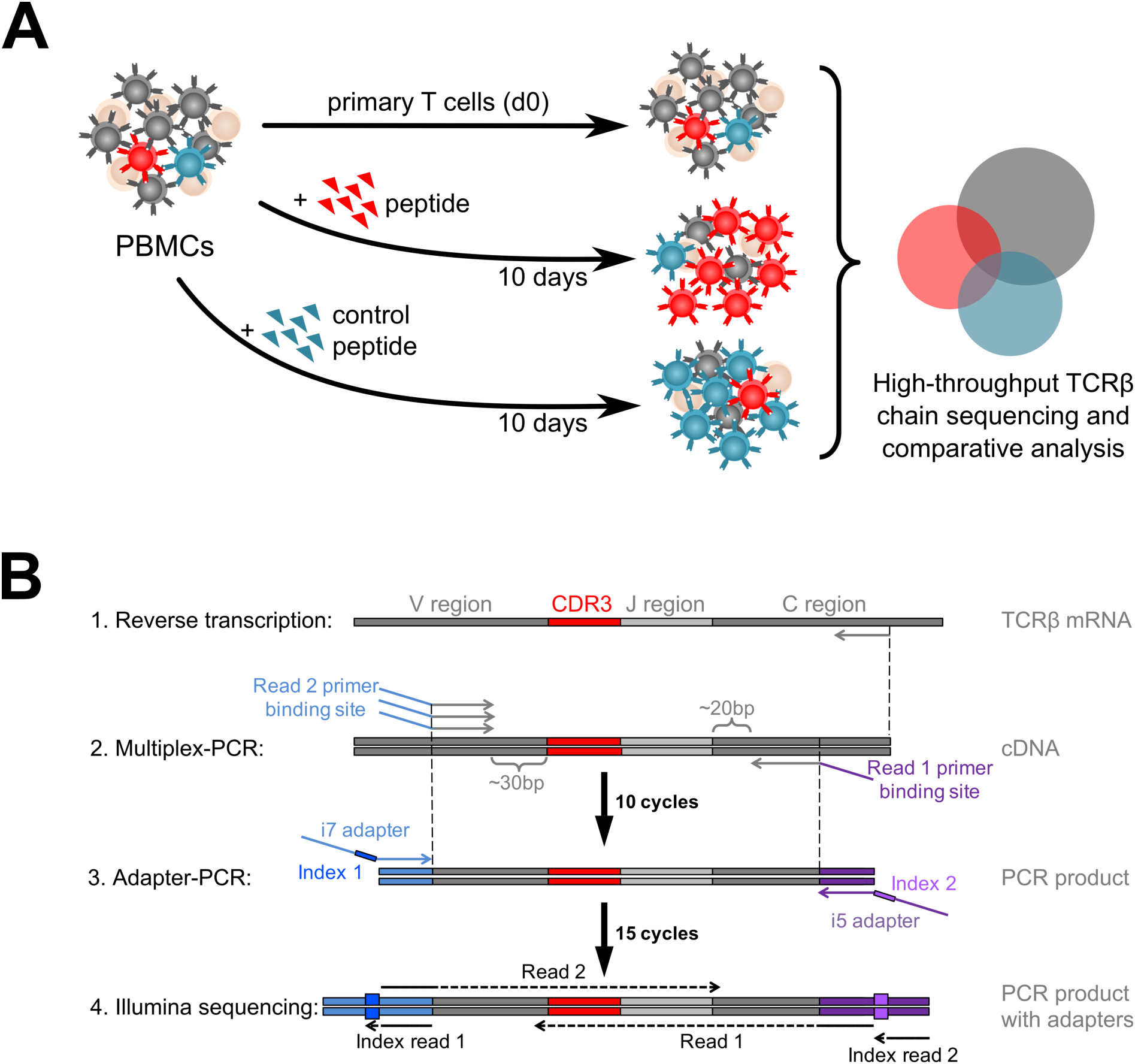
Experimental setup. (**A**) Schema of the three-sample assay used to expand and analyze peptide-specific T cells. PBMCs were isolated from peripheral blood of healthy donors, loaded with single CMV-derived peptides, and cultured for 10 days with IL-2. Cells before and after stimulation were lysed, and TCRβ libraries were prepared from bulk RNA and analyzed by high-throughput sequencing. Specific TCRβ sequences for each epitope were identified by comparing TCRβ clonotype frequencies in three samples, (i) stimulated with specific peptide, (ii) stimulated with control peptide, and (iii) before stimulation. Clonotypes that were enriched in condition (i), but not in controls (ii) and (iii), were considered specific. (**B**) Preparation of TCRβ libraries for bidirectional sequencing of the CDR3. After total RNA isolation, TCRβ RNA was reversely transcribed using a Cβ gene-specific primer. In a first PCR step, cDNA was amplified by semi-multiplexed PCR with a mix of 42 forward primers that covered all Vβ genes and appended the Illumina sequencing read 2 primer binding site to the product, and a reverse primer that was complementary to both Cβ genes and appended the sequencing read 1 priming site. A second PCR step was performed with a single primer on each side which adds Illumina i5 and i7 adapters and sample indices for multiplexing.

### High-throughput sequencing and data analysis

The barcoded samples were combined to a final concentration of 10 nM DNA and sequenced with the Illumina HiSeq1500 system in paired-end rapid run mode. The libraries were bidirectionally sequenced with read lengths between 120–175 bp in each direction.

Raw data were demultiplexed and quality-filtered using web-based tools on the Galaxy platform. Next, all reads were aligned to the matching Vβ, Jβ and Cβ genes, TCRβ clonotypes were built from identical sequences, and similar clonotypes were clustered with the MiXCR software (38). TCR clonotype data were further processed using custom scripts in R, in order to compare samples and characterize specificity and sharing of TCRβ sequences. Graphs were made with R, GraphPad Prism, or Microsoft Excel. P values were calculated with two-sided Mann-Whitney U tests in GraphPad Prism, version 7.

### Identification of specific TCRβ sequences

Specific TCRβ clonotypes for each epitope were identified by comparing TCRβ clonotype frequencies in three samples: one stimulated with the specific peptide of interest (S), one stimulated with a control peptide (C), and unstimulated PBMCs (U). Specific TCRβ clonotypes were required to be enriched in S over C and in S over U. Let s_i_, c_i_ and u_i_ be the relative read frequency (proportion of reads) of clonotype i in the three samples. To count as specific, clonotypes must exceed two enrichment cut-offs (Supplementary Figure 1, panel B). The first enrichment cut-off was identified as a local minimum of a weighted density distribution of log_10_ (s_i_/c_i_) of all medium- to high-frequency clonotypes, i.e. all clonotypes i that fulfilled the condition s_i_c_i_ > 10^−6^. Analogously, the second enrichment cut-off was identified as a local minimum of a weighted density distribution of log_10_ (s_i_/u_i_) of all clonotypes i that fulfilled s_i_u_i_ > 10^−7^. To eliminate low-fidelity background signals, specific clonotypes must also exceed a specific sample read count cut-off (Supplementary Figure 1, panel C). This cut-off was determined by analyzing the two density distributions of log_10_ s_i_ for all clonotypes i that had a low frequency in control samples, i. e. an absolute frequency of 1 to 10 reads in samples C or U, respectively. The read count at a local minimum of each of these two distributions was identified, and the mean of these two read counts served as the cut-off value. If any of these two density distributions did not have a local minimum, the cut-off was positioned at its global maximum times 100, i.e. at 100 reads in sample S or higher.

Using these criteria, specific TCRβ clonotypes were identified for 8 CMV-positive donors (P01-P08) and four epitopes (CRV, FRC, RPH, and TPR), resulting in identification of 1052 CMV peptide-specific TCRβ sequences which were unique at the amino acid level (all specific sequences are listed in Supplementary Table 2). Identification of antigen-specific TCRβ sequences from mini-LCL stimulations was achieved in a similar manner by comparing the frequencies of each clonotype in the CMV antigen-stimulated sample to their *ex vivo* frequencies and the frequencies in the samples obtained by stimulation with an empty-vector mini-LCL control. Only TCRβ sequences that were enriched compared to both control samples were considered specific for epitopes from that CMV antigen.

### Identification of public and related TCRβ sequences

Public CMV epitope-specific TCRβ sequences and TCRβ families were identified based on 1052 specific TCRβ sequences from donors P01–P08. In a first step, TCRβ sequences were categorized as public if they were CMV-specific in at least two donors with identical Vβ and Jβ gene segments, CDR3 amino acid sequence, and epitope specificity. In a second step, this set of sequences was extended by such TCRβ sequences that were present in only one donor, but highly similar to public TCRβ sequences. Sequences were considered highly similar if (a) they used the same Vβ gene segments, (b) had the same CDR3 length, and (c) differed in maximally two amino acids in the CDR3.

## Results

### A method for high-resolution TCRβ repertoire analysis by peptide stimulation

We devised a method to analyze human epitope-specific TCRβ repertoires at high resolution. This method combines simple short-term *in vitro* stimulation of peripheral blood mononuclear cells (PBMCs) with synthetic peptide (39, 40) and high-throughput TCRβ sequencing (Figure 1). Our focus was on donors carrying the HLA class I haplotype B*07:02∼C*07:02, which is the most frequent HLA-B/C haplotype in donors of European descent (41). We tested the four most immunogenic CMV peptides that are known to be presented by HLA allotypes encoded by this haplotype (17, 19, 42): the HLA-C*07:02-restricted epitopes CRVLCCYVL (CRV) and FRCPRRFCF (FRC) and the HLA-B*07:02-restricted epitopes RPHERNGFTVL (RPH) and TPRVTGGGAM (TPR). HLA-C*07:02-restricted epitopes are of special clinical interest because their recognition resists viral immunoevasion (17, 18). HLA-C*07:02 is prevalent not only in people of European descent, but also in East Asian and Native American populations (41).

TCRβ cDNA libraries for Illumina sequencing were prepared from T-cell samples in a 2-step RT-PCR (Figure 1B). The 2-step PCR procedure was designed to limit potential amplification bias due to multiplex priming and to increase fidelity by enabling bidirectional sequencing of the CDR3. After Illumina sequencing, TCRβ clonotypes were built using the software MiXCR (38). A median of 5.0 × 10^6^ productive TCRβ sequence reads were obtained per sample (Supplementary Table 2). When we plotted TCRβ clonotype frequencies before and after stimulation, well-separated clusters suggestive of peptide-reactive expanded TCRβ clonotypes became apparent in CMV-positive, but not in CMV-negative donors (Figure 2). For a more precise definition, we evaluated TCRβ sequences to be epitope-specific if they appeared in the cluster of enriched clonotypes after cultivation with a specific peptide, but not after cultivation with a control peptide of different HLA restriction (three-sample comparison, enrichment cut-off). In addition, a frequency cut-off was applied to minimize statistical noise from low-frequency clonotypes (Supplementary Figure 1); this cut-off was calculated from the sample-specific clonotype frequency distribution.

**Figure 2.**
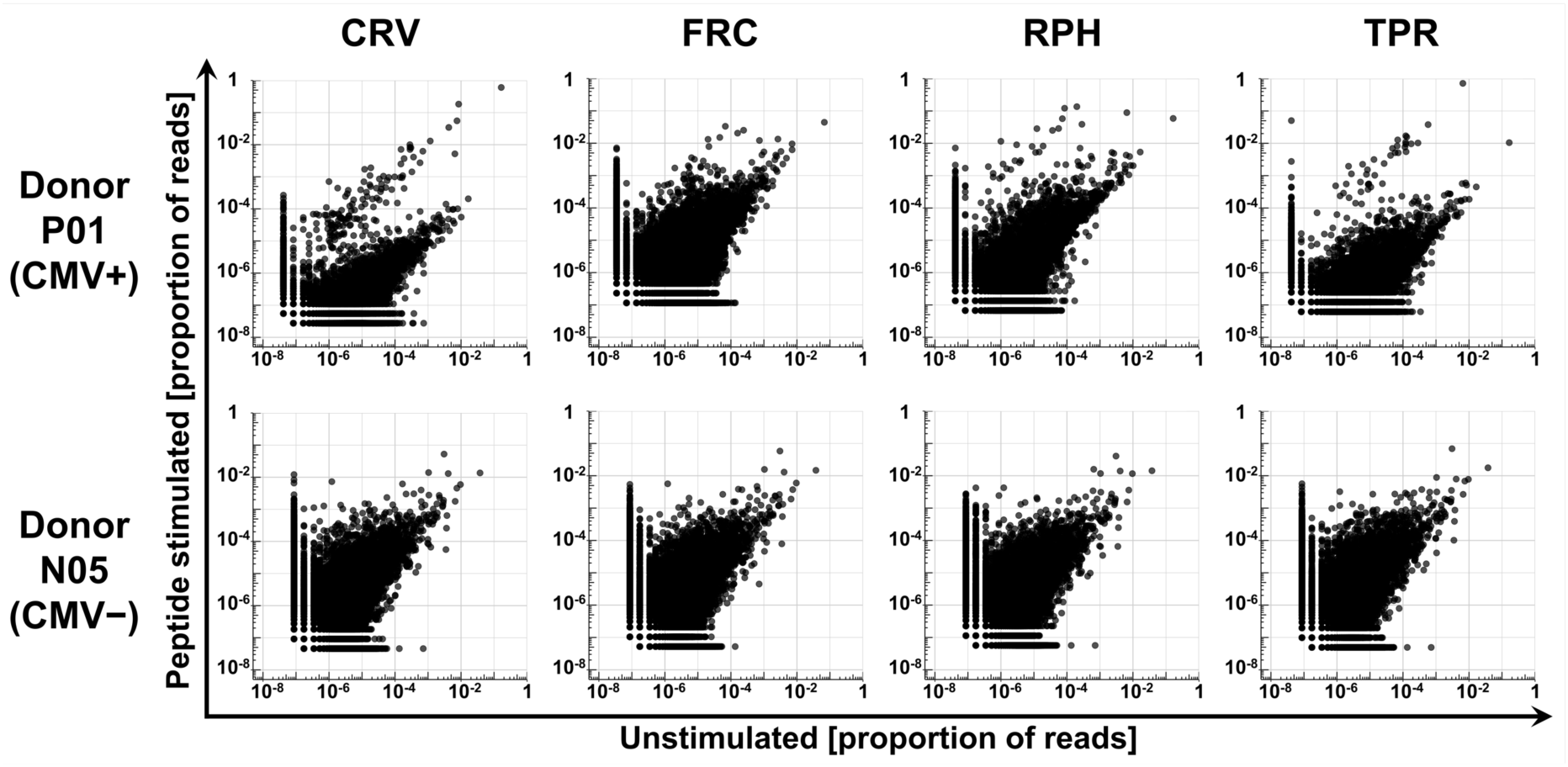
Populations of enriched TCRβ clonotypes are exclusive to CMV positive donors. Relative frequency (proportion of reads) of TCRβ clonotypes before (x-axis) and after (y-axis) stimulation with one of four CMV peptides in CMV-positive donor P01 (upper panel) and CMV-negative donor N05 (lower panel). Each TCRβ clonotype is defined as the entirety of identical reads on the nucleotide level and represented by a black dot. Clonotypes that were undetectable in one condition were assigned a pseudofrequency corresponding to 0.5 reads to enable their display on a logarithmic axis.

We identified a total of 1052 unique TCRβ amino acid sequences in 8 CMV-positive donors (P01–P08) that met the specificity criteria for exactly one epitope (listed in Supplementary Table 2). Of these, 435 were specific for CRV, 266 for FRC, 191 for RPH, and 160 for TPR. Nineteen TCRβ sequences passed the criteria for two epitopes, no TCRβ for more than two epitopes. Thus, we observed minor overlap between TCRβ sequences assigned to different specificities.

### Peptide stimulation expands T-cell clonotypes that recognize processed antigen

There is a concern that synthetic peptide may not exclusively stimulate T cells that will be able to recognize the naturally processed epitope (43). Therefore, we tested whether our peptide-enriched T-cell clonotypes respond to endogenously processed CMV antigens. We established autologous mini-lymphoblastoid cell lines (mini-LCLs) that constitutively express CMV proteins pp65 or IE-1 from a mini-Epstein-Barr virus genome. Such mini-LCLs effectively present CMV epitopes of any autologous HLA restriction to CD8+ and CD4+ T cells (17, 37, 44, 45). PBMCs of 3 CMV-positive donors, P01-P03, were stimulated with autologous mini-LCLs (pp65, IE-1, or control mini-LCLs without CMV antigen), and the resulting TCRβ repertoires (Supplementary Table 2) were compared with those obtained through peptide stimulation. In the mini-LCL condition, TCRβ sequences were considered CMV-specific if they were enriched by stimulation with mini-LCLs that expressed the CMV antigen of interest, but not by stimulation with control mini-LCLs that lacked the CMV antigen. We found that the majority of TCRβ clonotypes that were expanded by one of the peptides CRV, RPH, or TPR also recognized the corresponding CMV antigen processed by mini-LCLs (Figure 3). These results confirm that our simple peptide stimulation assay is a generally valid approach for the identification of CMV-specific TCRβ clonotypes of both high and low frequency.

**Figure 3.**
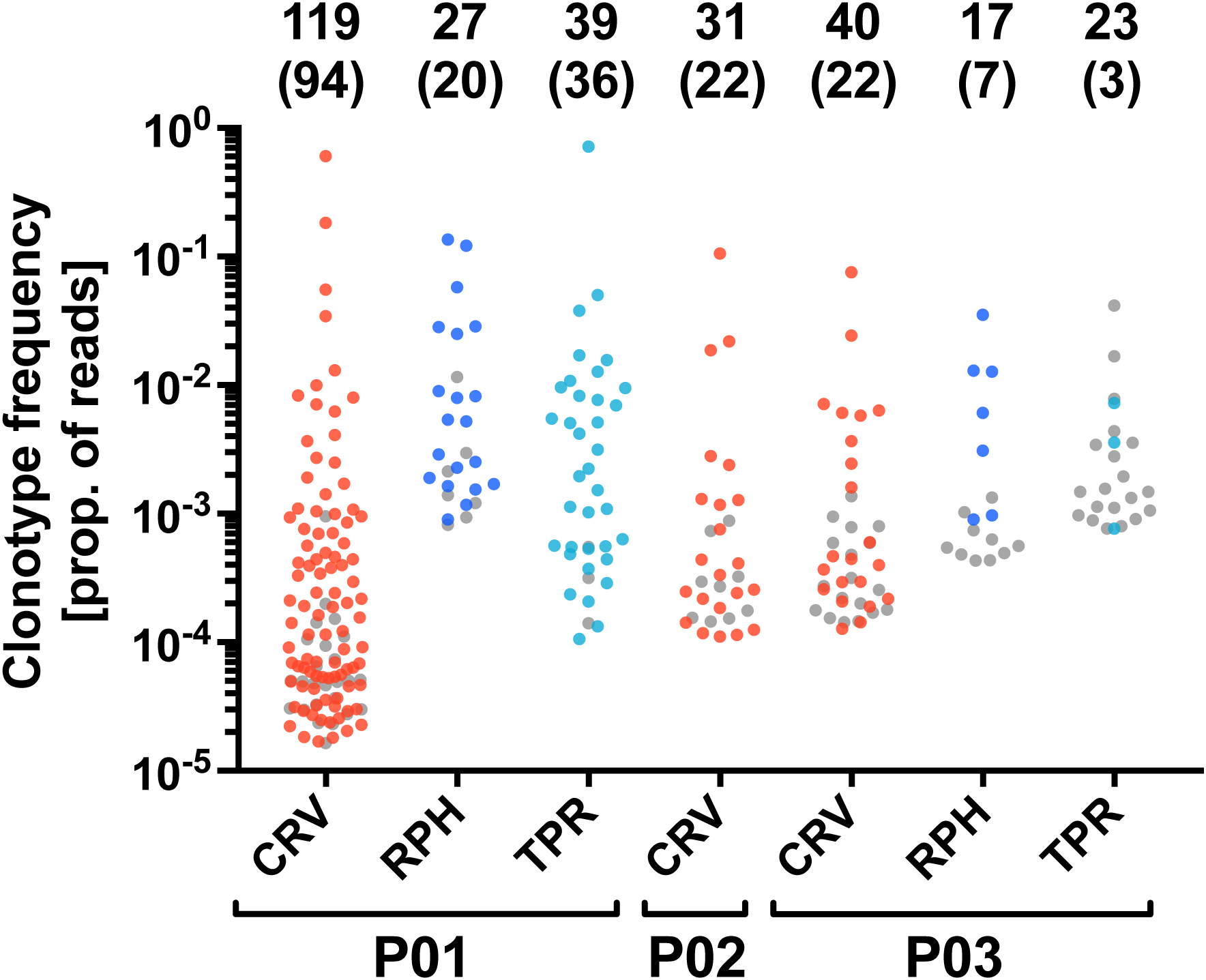
Specific TCRβ clonotypes identified in the peptide stimulation assay also respond to endogenously processed antigen. The analysis was performed for the IE-1 antigen (epitope CRV) in CMV-positive donors P01, P02, and P03, and for the pp65 antigen (epitopes RPH and TPR) in donors P01 and P03. The plot shows all TCRβ clonotypes that were identified as epitope-specific in the peptide stimulation assay. The y-axis indicates the frequency of each clonotype after peptide stimulation. Colored dots represent clonotypes that were specifically enriched by stimulation with an autologous mini-LCL that expresses the corresponding CMV antigen, grey dots indicate clonotypes for which this was not the case. The numbers on top indicate the total number of epitope-specific TCRβ clonotypes and, in parentheses, the number of clonotypes responding to antigen endogenously processed by mini-LCLs. Samples of donor P01 were CD8-enriched before sequencing.

### CMV-specific TCRβ clonotypes are abundant in the T-cell repertoire of virus carriers

We investigated the contribution of CMV epitope-specific T-cell receptors to the overall TCRβ repertoire in peripheral blood of 8 CMV-positive (P01-P08) and 8 CMV-negative (N01-N08) donors. Among the top 100 most frequent *ex vivo* TCRβ clonotypes of any CMV-positive donor (Figure 4A), 2 to 10 were specific for one of the CMV epitopes CRV, FRC, or TPR. In 7 of 8 CMV-positive donors, CMV-specific clonotypes were among the 5 most frequent clonotypes, and in 2 of 8 donors, they supplied the top-frequency clonotype of the entire repertoire. Among a donor’s CMV-specific clonotypes, the most frequent one was specific for CRV in 6 and specific for FRC in 2 of 8 donors. In CMV-negative donors, none of the top 100 TCRβ clonotypes were specifically enriched by CMV peptide stimulation, and thus none of them was categorized as CMV-specific. When looking at the cumulative read frequencies, CRV- or FRC-specific clonotypes dominated the response in CMV carriers (Figure 4B). These results demonstrate that CMV-specific T cells distinctly shape the T-cell repertoire of virus carriers, with a prominent role for HLA-C-restricted clonotypes.

**Figure 4.**
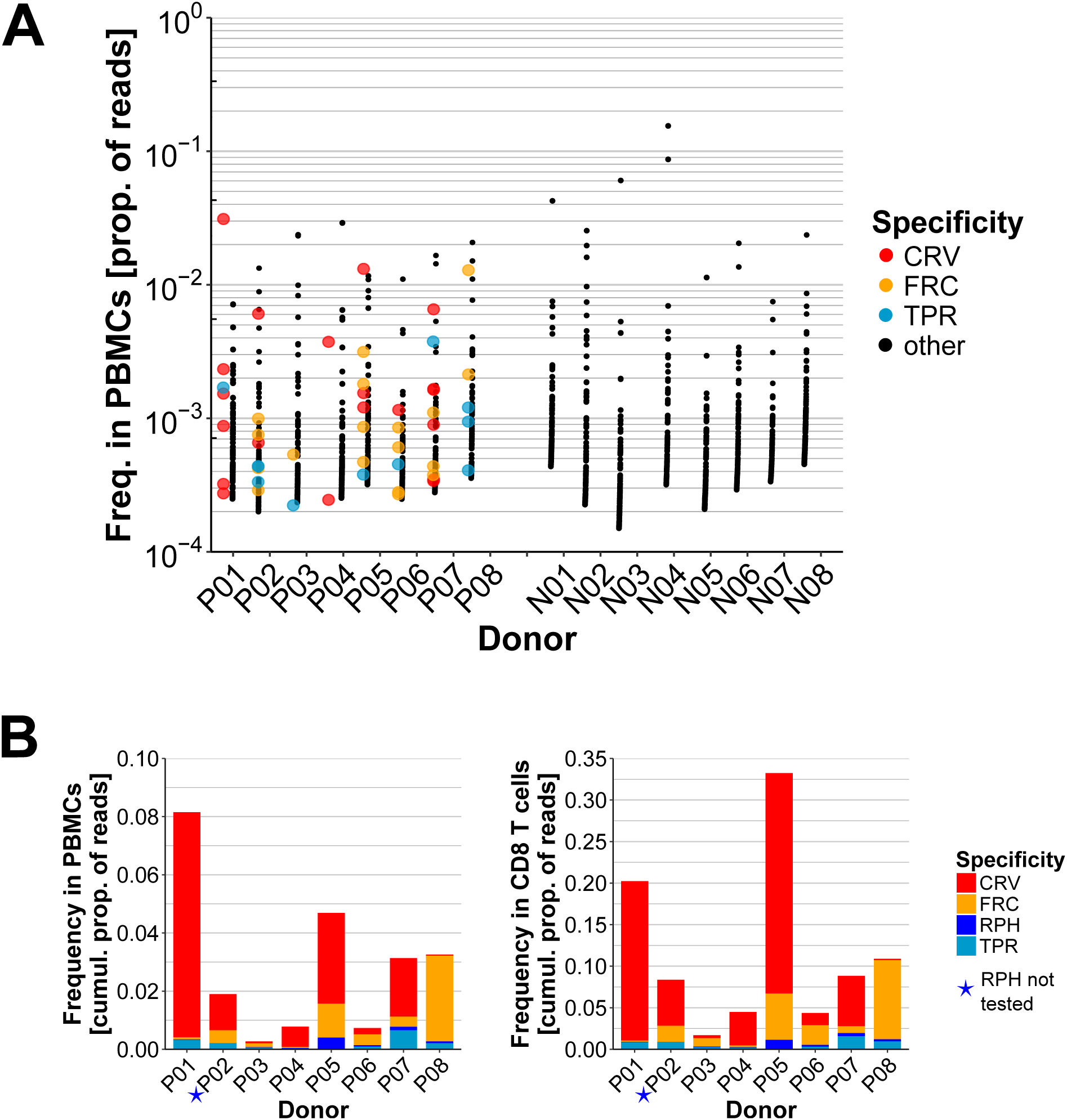
CMV-specific TCRβ clonotypes dominate the peripheral T-cell repertoire of CMV-positive donors. (**A**) The figure shows the proportion of reads of the 100 most frequent TCRβ clonotypes of CMV-positive donors P01-P08 and CMV-negative donors N01-N08 in peripheral blood *ex vivo*. TCRβ clonotypes that were identified as specific for CMV epitopes CRV, FRC, or TPR are shown as colored dots, clonotypes of unknown specificity as black dots. Because epitope RPH was not tested in donors P02 and N01-N08, it was omitted from this analysis. (**B**) Frequencies of CMV-specific TCRβ sequences in the *ex vivo* repertoires (left) and CD8-enriched repertoires (right) of CMV-positive donors P01-P08 as cumulative proportion of reads. RPH-specific T cells were not investigated in donor P02 and such T cells are therefore not depicted in the plots.

### Patterns of Vβ and Jβ gene segment usage in CMV epitope-specific TCRs

We analyzed TCRβ clonotypes sharing their epitope specificity for shared structural features. First, we evaluated overall use of Vβ and Jβ gene segments (Figure 5) in CMV-specific TCRβ clonotypes. For each epitope, particular Vβ and Jβ genes were overrepresented, such as Vβ-6-1/-5/-6, Vβ25-1 and Vβ28, and Jβ1-1, Jβ2-1 or Jβ2-7 for epitope CRV. However, no Vβ-Jβ combination dominated the response to any of the four epitopes. It follows that gene segment use alone is not sufficiently informative as a marker of CMV-specific CD8+ T-cell immunity to the epitopes studied here.

**Figure 5.**
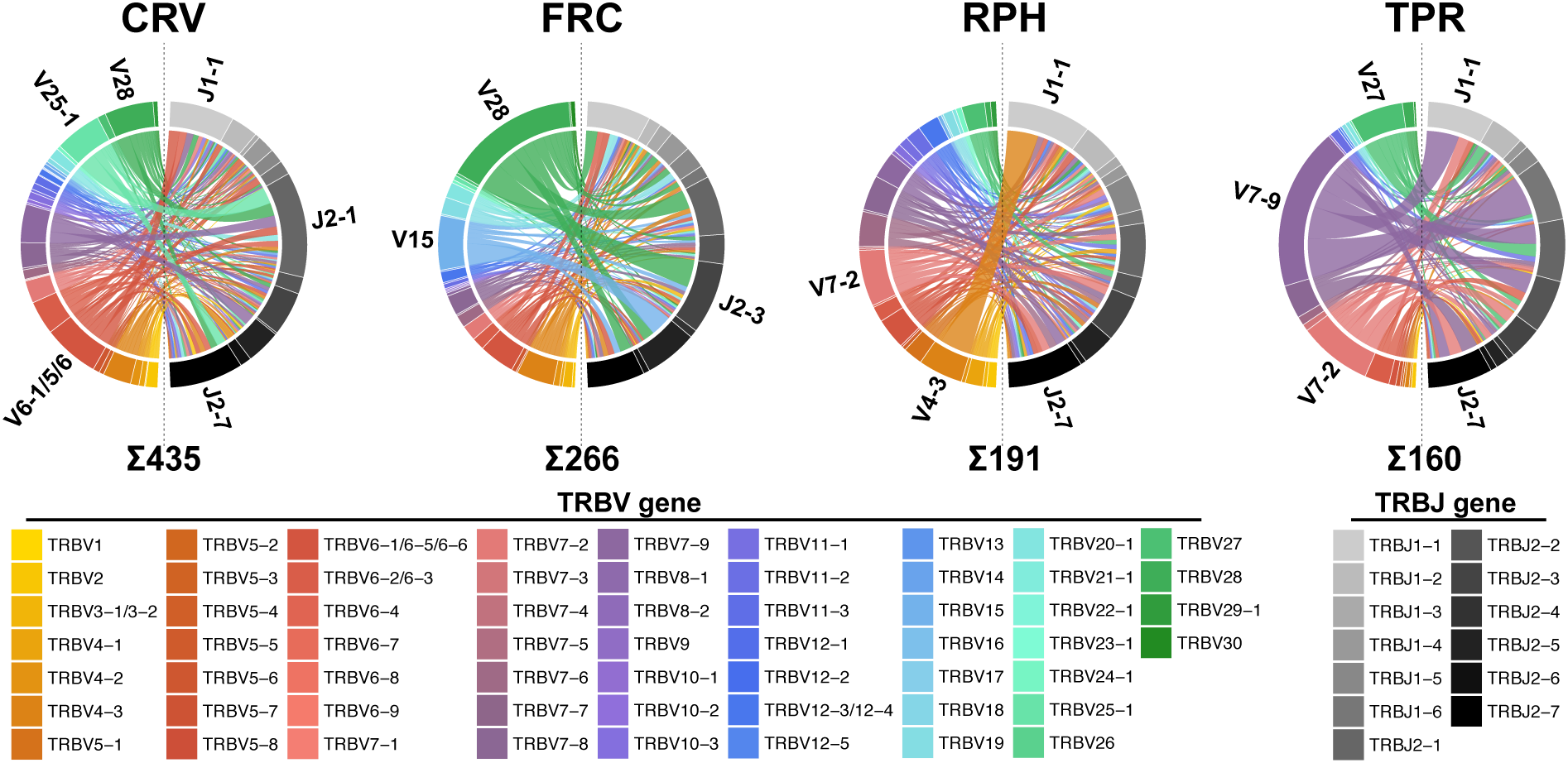
Usage of Vβ and Jβ gene segments in CMV epitope-specific T cells. The left semicircle of each chord diagram represents Vβ gene segment usage (rainbow colors), the right semicircle represents Jβ usage (grayscale shades), and the chords indicate which gene segments appear together in TCRβ sequences. Sizes of sectors and chords are proportional to the sum of the number of nucleotide-unique antigen-specific TCRβ sequences in each of the donors P01-P08. The most frequently used gene segments are labeled and the numbers below the plots show how many TCRβ sequences were specific for each epitope.

### Public CMV-specific TCRβ sequences and TCRβ families

Next, we searched for the presence of identical TCRβ amino acid sequences with the same CMV epitope specificity in different donors, known as shared or public TCRs. We found 26 TCRβ sequences that were specific in at least 2 out of 8 CMV-positive donors (Table 1). Several of these public sequences with the same epitope specificity were closely related in sequence, since they used the same Vβ gene, had the same CDR3 length, and differed in a maximum of 2 amino acids within the CDR3. Hence, we looked for additional TCRβ sequences that appeared in only one of the 8 donors, but were closely related to one of the 26 public TCRβ sequences according to the criteria stated above; we identified 37 such TCRβ sequences. The resulting set of 63 public or related CMV-specific TCRβ sequences was composed of 17 similarity groups, which we refer to as public TCRβ families. Of the 63 sequences, 21 were HLA-B-restricted (epitopes TPR and RPH), and 10 of them had been previously described (29, 42, 46, 47). In contrast, the epitope specificity and HLA restriction of none of the 42 HLA-C-restricted TCRβ sequences (epitopes CRV or FRC) had been previously shown, although 5 of these sequences were found to be enriched in CMV-positive donors compared to CMV-negative donors (48).

**Table 1.**
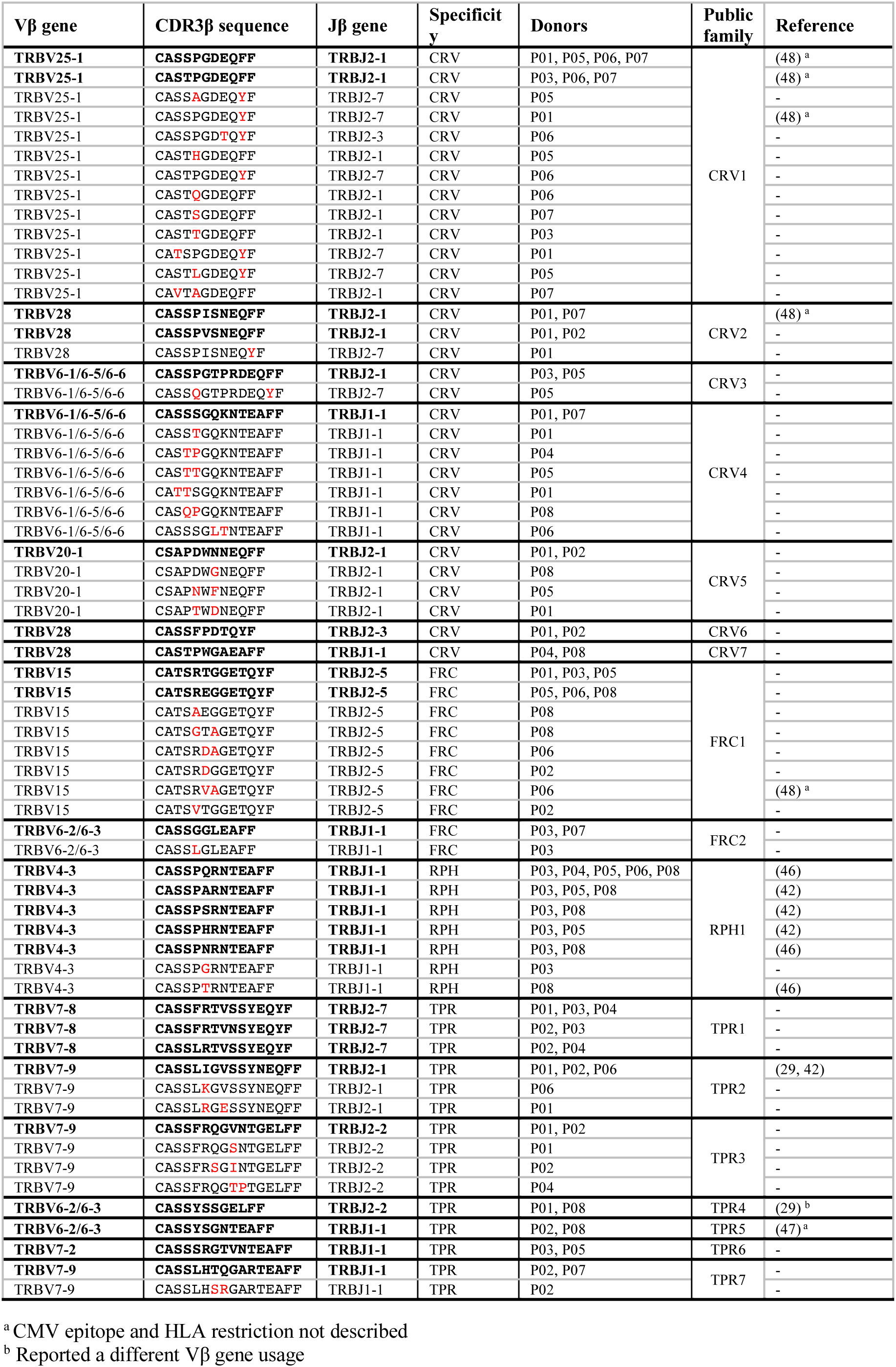
The 63 public or related TCRβ sequences identified in this study. Only donors are listed in whom the TCRs were functionally identified as epitope-specific. The 26 public TCRβ sequences are in boldface. Amino acid exchanges in the 37 related TCRβ sequences compared to the most similar public TCRβ sequence are highlighted in red.

### Public TCRβ families precisely distinguish CMV-positive from CMV-negative donors

We analyzed the frequency of public TCRβ families in primary T cells from an extended cohort of CMV-positive and CMV-negative donors: 8 additional CMV-positive donors (P09-P16) and 9 CMV-negative donors (N01-N09) with the HLA-B*07:02∼C*07:02 haplotype. Public TCRβ family members were much more abundant in CMV-positive than CMV-negative donors (Figure 6A). This was true for each of the four CMV epitopes when evaluated separately (Figure 6B, left). Taking all epitopes together, the median cumulative read frequency was 171-fold higher in CMV-positive than in CMV-negative donors of our 25-donor cohort (Figure 6C; data for each TCRβ and donor are provided in Supplementary Table 3). Mean cumulative frequency in donors P09-P16 and donors P01-P08 was similar, which showed that there was no bias in favor of the P01-P08 cohort of donors in whom the sequences were originally identified. To further validate the set of public TCRβ families, we tested it on a larger cohort of donors whose *ex vivo* TCRβ repertoires were recently published (48) and whose CMV status and partial HLA type (low resolution HLA-A and -B, no HLA-C) was available. Since presence of HLA-B7 is a strong indicator of the presence of the haplotype HLA-B*07:02∼C*07:02 in persons of European descent, whereas persons of Asian descent often express HLA-C*07:02 without HLA-B*07:02 (41), we limited our analysis to the 352 donors categorized as “White, not Hispanic or Latino”. Of these donors, 94 were HLA-B7-positive, and 258 were HLA-B7-negative. As shown in Figure 6C, CMV-specific public TCRβ families were strongly enriched in HLA-B7-positive, CMV-positive donors, but not in HLA-B7-positive, CMV-negative donors (P < 1 × 10^−15^). In HLA-B7-negative donors, no enrichment was observed irrespective of CMV status. In a separate analysis of each epitope in this cohort, CRV was the strongest discriminator (P = 7.5 × 10^−13^; Figure 6B, right).

**Figure 6.**
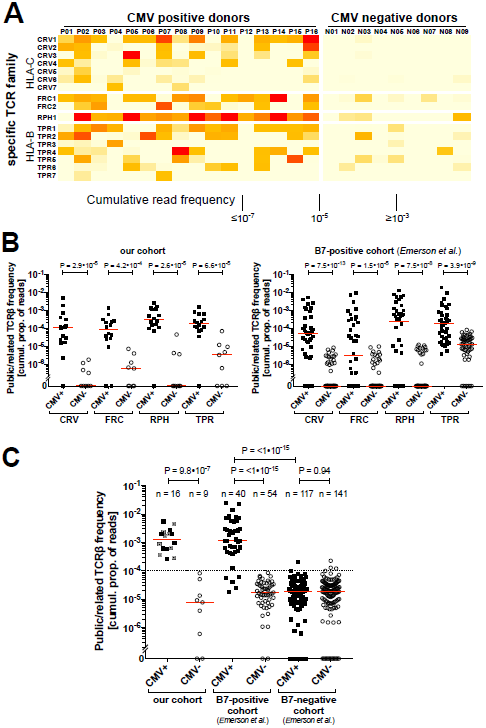
Frequency of public TCRβ families in peripheral T-cell repertoires of CMV-positive and CMV-negative donors. (**A**) Cumulative proportion of TCRβ sequence reads for each individual TCRβ family in the *ex vivo* TCRβ repertoires of donors P01-P16 and N01-N09. Read frequencies for each of the 63 TCRβ sequences that form these families can be found in Supplementary Table 3. (**B**) Cumulative proportion of reads for all public or related TCRβ sequences with the same epitope specificity in our donor cohort and an independent HLA-B7-positive subcohort from Emerson et al. (48) (**C**) Cumulative proportion of TCRβ sequence reads of all public/related TCRβ sequences in CMV-positive (solid circles) and CMV-negative (hollow circles) donors in our cohort, and in HLA-B7-positive and HLA-B7-negative donors from Emerson et al. (Supplementary Table 3). Grey circles identify donors P01-P08 from whose repertoires the TCRβ sequences were originally derived. The dashed line indicates a possible cut-off to separate CMV-positive and CMV-negative donors in our cohort (F1 score = 1) and the B7-positive cohort of Emerson et al. (F1 score = 0.93). Red lines show the median cumulative read frequencies. P values were calculated with a two-tailed Mann-Whitney U test.

A chosen cut-off value of 10^−4^ for the total proportion of reads of our set of 63 public or related TCRβ sequences lead to perfect discrimination between CMV-positive and CMV-negative donors within our cohort, and very good identification of CMV-positive donors within the HLA-B7-positive published cohort (F1 score = 0.93; 100% specificity, 88% sensitivity). Taken together, our results show that the CMV-specific TCRβ signature, as identified through our approach, is highly indicative of CMV-specific T-cell immunity associated with the CMV status in healthy donors.

## Discussion

Here we analyzed the composition and sharing of TCRβ repertoires against four epitopes that are major targets of the CMV-specific CD8+ T-cell response. Specific TCRs against two of these epitopes, restricted through HLA-C, had not been studied before. We identified a set of CMV-specific public TCRβ families that distinguishes CMV-positive from CMV-negative healthy persons in two independent cohorts with high precision.

We show that the CMV epitopes studied here considerably shape the T-cell repertoire of healthy virus carriers. HLA-C*07:02-restricted T cells were particularly prominent; in 6 of 8 donors they provided one of the four most frequent TCRβ clonotypes to the overall T-cell repertoire, and in 2 of these donors the top-frequency TCRβ clonotype. These results expand on the general observation that CMV-specific T cells make up for a large proportion of the CD8+ T-cell repertoire, on average amounting to 5% of the total CD8+ T-cell response based on interferon-γ secretion (49). This proportion is likely to be even larger when all antigen-specific cells are included in measurements, not just those that exert a chosen effector function at the time of analysis (50). Advanced donor age boosts CMV-specific T-cell frequencies as well (19), an aspect we could not investigate in our cohort of anonymous donors.

The TCRβ signatures of the CMV epitope-specific T cells studied here were diverse. For a given epitope, the dominant TCRβ sequence was usually a different one in different donors. Thus, TCRβ repertoires were not dominated across the board by heavily conserved (public) sequences, in contrast to what was observed for some epitopes from influenza (51, 52) or EBV (53). Rather, the TCRβ repertoires studied here were mostly composed of clonotypes that were shared with only a subset of matched donors, or were entirely donor-specific (private). These patterns are reminiscent of those previously found for TCR repertoires of HLA-A- and HLA-B-restricted CMV-specific T cells (29–31, 33, 34, 54, 55). Nonetheless, our approach identified a series of public TCRβ chains whose cumulative frequency was strongly indicative of CMV carrier status in larger cohorts of healthy donors, even though these public TCRs did not necessarily represent the most abundant epitope-specific T cells in the 8 donors in whom they were originally identified. It seems clear that exclusive TCRβ sequencing underestimates the true diversity of an epitope-specific T-cell response, since the same TCRβ can be paired with different TCRα chains to generate the same or an overlapping human antiviral epitope specificity (53, 56, 57). However, our results show that TCRβ sequencing, even in the absence of TCRα analysis, is already highly informative regarding the CMV-specific T-cell status of donors. This finding may already have been anticipated by a recent large-scale study (48), which showed that signatures of enriched TCRβ sequences distinguished CMV-positive from CMV-negative donors with high precision. Our study extends these findings by demonstrating the predictive power of TCRβ sequences with known CMV epitope specificity. In contrast, the previous study (48) analyzed TCRβ sequences that were CMV-associated, but mostly not known to be CMV-specific, which means that other pathogens with overlapping epidemiology may have contributed to the signal. For future diagnostic or prognostic applications in immunocompromised patients, it will be preferable to focus on analysis of precisely defined antigen-specific TCRβs, since such patients may simultaneously reactivate or acquire multiple, and even related, pathogens. We conclude that TCRβ sequencing provides a highly informative and economic standalone approach to identification of epitope-specific T cells in healthy carriers and patients at risk of viral reactivation and in potential need of antiviral prophylaxis or treatment (1, 58, 59).

Conserved TCRβ chains are not necessarily perfectly conserved. It was often observed that human HLA-A- or HLA-B-restricted CD8+ T-cell responses to an epitope contain non-identical but highly similar TCRβ chains; these use the same or closely related Vβ and Jβ segments and typically have CDR3 regions that show exchanges in only few amino acid positions. Such relationships are apparent in datasets on multiple epitopes from CMV (29, 31, 42, 55, 60, 61), EBV (29, 31, 53, 61–64), HIV (29), or influenza virus (51, 52). Similarity between TCRβ chains of the same specificity can also manifest as the presence of conserved short motifs within the CDR3 (29, 52), often at or near residues 6 and 7; these residues are at its center and generally make strong contributions to binding of MHC/peptide by the TCR and shaping T-cell specificity (65). Analysis of such TCRβ relationships was recently taken to a more comprehensive level (33, 34) by the design of computational algorithms to cluster T cells of the same specificity according to multiple indicators of sequence similarity. Global sequence similarity of the CDR3 is the single factor that predicts specificity best (34), and prediction is further improved by adding aspects such as CDR3 length and Vβ gene segment usage. Accordingly, we observed multiple occurrences of high similarity of CDR3 sequence, length and Vβ usage among our 26 CMV-specific public TCRs. Therefore, we decided to group CMV-specific public and related TCRβs into families based on these criteria. We found that TCRβ families that were defined in this way could precisely distinguish healthy persons with and without established CMV-specific T-cell immunity. Public TCRβ family sequences were as rare in CMV-negative persons who carry the relevant HLA haplotype as in CMV-positive persons who lack the HLA haplotype. This finding shows that such TCRβ family members are unlikely to appear in T cells of irrelevant specificity or HLA restriction, confirms the predictive power of TCRβ sequencing, and suggests that our TCRβ grouping approach can in future studies be extended to a broader variety of epitopes, viruses, or antigenic entities.

Since its initial description (26, 32), high-throughput sequencing of TCRβ repertoires has become a widely used research method, but it has not yet found wide clinical application. We have now introduced technical improvements that will enhance its robustness and applicability to accelerate advancement to routine clinical use. Dual-indexed, paired-end bidirectional sequencing of the entire CDR3 region is likely to reduce errors in the resulting Illumina sequencing data (66). Our technique to identify epitope-specific TCRβ repertoires by peptide stimulation and comparison to two controls eliminates the need to physically isolate specific T cells. Earlier studies that aimed at characterizing epitope-specific TCR repertoires have generally employed sorting of HLA/peptide multimer-labeled T cells (33–35, 54, 61), or occasionally sorting of T cells labeled with markers of activation or proliferation (67). High-throughput sequencing has the capacity to identify TCR sequences even from very low-frequency components of the sample. However, T-cell subsets sorted based on multimers or other markers are not perfectly pure, and there is a possibility that low-frequency contaminants, which may derive from dominant clones from the parent population, are erroneously assigned the specificity of interest. Regardless whether multimer sorting or peptide stimulation is used to identify specific TCR clonotypes, artifacts can be avoided by quantitatively verifying enrichment of clonotypes relative to the parent population and relative to a sample treated with a different multimer or peptide. Our present approach has the limitation that it only covers T cells capable of proliferating *in vitro* in response to antigen. However, depending on the research question at hand, this limitation may be advantageous, since T cells capable of proliferation will in many cases be those that are functionally more relevant in disease or immune control. Moreover, the numerical increase of antigen-responsive T cells due to proliferation, as well as the increased absolute amount of TCRβ mRNA per cell several days after activation (68), will in itself increase resolution and thus the likelihood that rare clonotypes can be detected in samples of limited size. In contrast, HLA/peptide multimer staining can capture T cells that express a specific TCR irrespective of their functional properties; however, in spite of recent progress (69, 70), multimers are still challenging to produce for certain HLA allotypes and epitopes, and they may not always stain the entire antigen-specific T-cell population (71).

It seems safe to predict that identification and quantification of antigen-specific TCRβ repertoires will increasingly enter clinical practice for purposes of diagnostics and monitoring. Repositories of annotated antigen-specific TCRs (72), refined computational tools for TCR sequence analysis (38) and TCR sequence datasets (48), generously made available to the public, will be of great use in further developing the method. With growing datasets, an increasing number of epitope-specific TCR sequences against various pathogens will be found to be shared between carriers. Such public TCR sequences will find application in various fields. In clinical research, public TCR sequences may be used as indicators to track virus-specific T-cell responses in patients after transplantation in order to identify epitopes that mobilize a protective T-cell response against pathogens such as CMV (6). Virus-specific TCR signatures may also be exploited in diagnostics and disease monitoring (36, 73, 74) to inform about a patient’s status regarding past or present infection with multiple pathogens, success of vaccination or T-cell transfer, and risk of future infection or reactivation. Pathogen-specific TCRs that are frequently present in the self-tolerant repertoire of multiple healthy donors are likely to be non-responsive to human antigens in various genetic backgrounds. Such TCRs carry a low risk of allo-HLA cross-reactivity (75) and are therefore favorable candidates for immunotherapy with TCR-transgenic T cells (76).

## Supplementary Material

**Supplementary Figure 1.** Criteria for identification of CMV-specific TCRβ clonotypes.

**Supplementary Table 1.** Donors, epitopes, and primers.

**Supplementary Table 2.** Sample metadata and specific TCRβ sequences.

**Supplementary Table 3.** Frequencies of 63 public or related TCRβ sequences in *ex vivo* repertoires of two independent HLA-matched and one HLA-mismatched donor cohort.

**Supplementary Figure 1.**
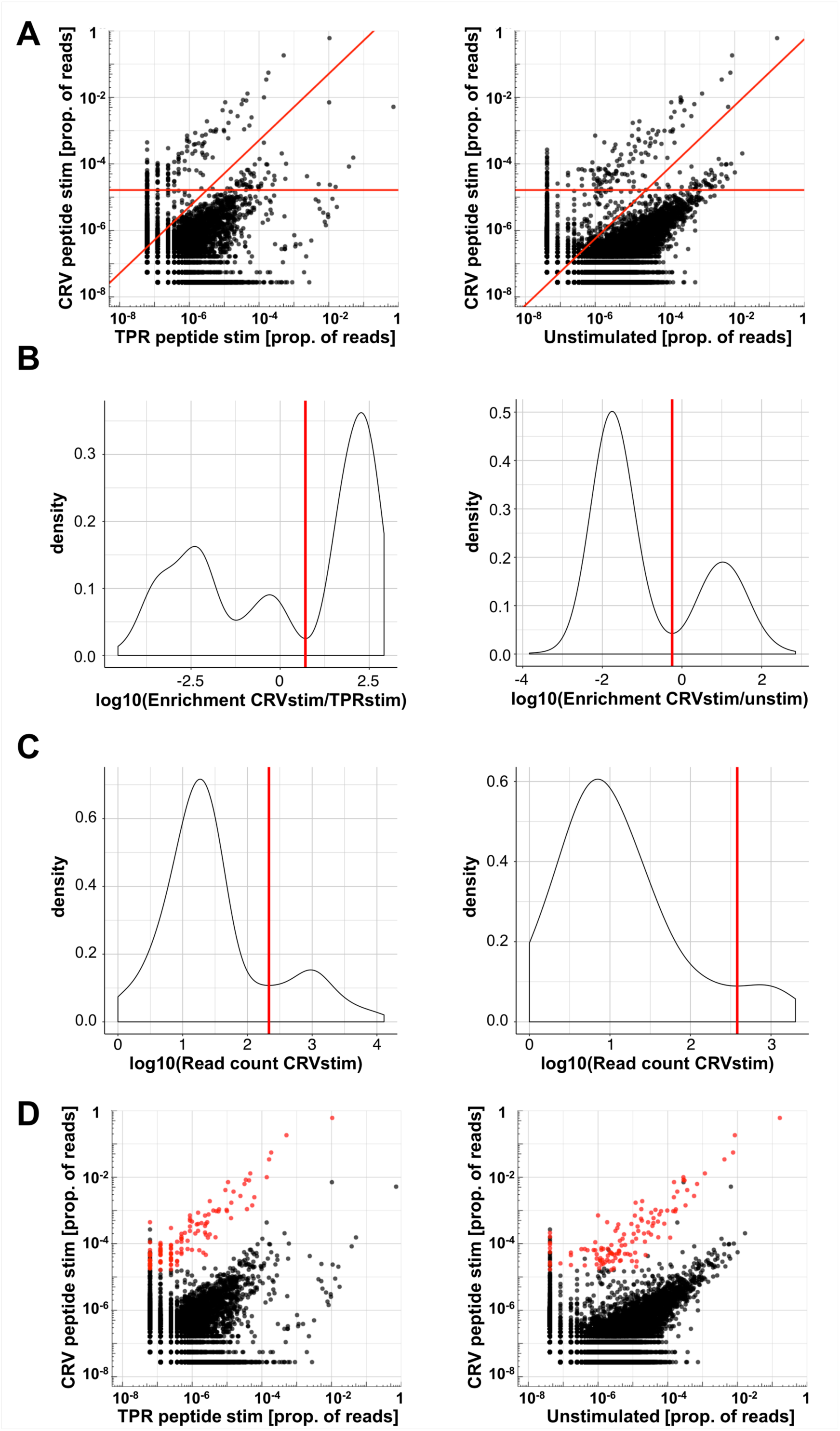
Criteria for identification of CMV-specific TCRβ clonotypes. As an example, representative data from donor P01 for identification of CRV-specific T cells are shown. Diagrams on the left relate to the binary comparison of the CRV-stimulated sample and the TPR-stimulated sample (control peptide). Diagrams on the right relate to the comparison of the CRV-stimulated sample and the unstimulated PBMC sample. Only TCRs that fulfilled all criteria were categorized as CRV epitope-specific. In dot plots, a frequency of zero is plotted as a pseudofrequency of 0.5 reads. (**A**) Relative frequencies of TCRβ clonotypes. Red lines indicate the enrichment cut-off (diagonal line) and the specific sample read count cutoff (horizontal line). (**B**) The enrichment cut-off (red line) is defined as the local minimum of the weighted density distribution of the logarithm of clonotype enrichment. Clonotype enrichment is the ratio of clonotype frequencies in the CRV-stimulated sample and either control. (**C**) Read count cut-offs (red line) were defined as local minima of the distribution of the logarithms of clonotype counts, determined for clonotypes with a read count of 1 to 10 in the TPR-stimulated (left) or unstimulated (right) sample. The mean of these two cut-offs was used as the final specific sample read count cut-off. (**D**) Relative frequencies of TCRβ clonotypes as in A, but TCRβ clonotypes that fulfil all inclusion criteria for being CRV epitope-specific are shown in red.

